# Structural basis of nonmuscle myosin-2 autoinhibition mechanisms

**DOI:** 10.1101/2025.10.23.684279

**Authors:** Sarah M. Heissler, Giovanna Grandinetti, James R. Sellers, Krishna Chinthalapudi

## Abstract

Nonmuscle myosin-2 (NM2) is a fundamental actin-based mechanochemical ATPase that regulates cellular architecture, migration, adhesion, and force generation across diverse biological contexts. NM2 function is tightly regulated by a structural transition between an autoinhibited monomeric (10S) conformation in which ATPase activity, actin binding, and filament assembly are coordinately suppressed and an enzymatically active, filamentous conformation. The autoinhibited conformation is critical for the spatial and temporal control of contractility in nonmuscle cells, yet structural insights into the 10S conformation remain largely elusive. Here, we present the cryo-EM structure of full-length human NM2B in the 10S conformation at ∼5.24 Å resolution. The structure reveals a tri-segmented tail fold that introduces asymmetric interactions between both myosin heavy chains and associated light chains and the sequestration of key interfaces required for actin binding and filament assembly. These structural insights provide a mechanistic foundation for understanding how regulatory post-translational modifications and disease-associated mutations shift NM2 conformational equilibria and may enable the development of structure-based interventions for cytoskeletal diseases including hearing loss, neurodegeneration, and cancer.

## Introduction

The ability of cells to dynamically alter their shape and mechanical properties in response to various stimuli is fundamentally dependent on cytoskeletal proteins of the myosin superfamily. One major protein that drives dynamic changes in the actin cytoskeleton through cellular force generation is nonmuscle myosin-2 (NM2), a ubiquitous and highly abundant actin-based mechanochemical ATPases in eukaryotic cells^1-3^. The NM2 heavy chain is composed of a motor domain and a tail domain connected by a neck domain that binds the myosin essential (ELC) and regulatory (RLC) light chain^4,5^. The motor together with the light chain bound neck domain is also referred to as the myosin head. The ∼ 1500 Å long tail is helical and terminates in a short non-helical tailpiece (NHT). Two copies of the heavy chain form a symmetric and extended coiled coil structure known as 6S monomer^6^. The motor domain binds filamentous actin and ATP and converts the chemical energy derived from the hydrolysis of ATP to mechanical force and movement of actin filaments^7^. NM2 functions predominantly as a ∼570 kDa asymmetric autoinhibited monomer known as 10S or as an active ∼16 MDa symmetric complex with 30 copies of the monomer assembled into a bipolar filament with a length of ∼300 nm^8,9^. A dynamic equilibrium between the autoinhibited monomer and the assembled, filamentous myosin pool allows nonmuscle cells to reorganize their cytoskeletons in response to regulatory inputs^3,10-15^. The equilibrium is precisely controlled by on-demand phosphorylation and dephosphorylation of the RLC that mediates the transition between the ‘off’ and the ‘on’ states, respectively^16,17^. The binary switch mechanism between a regulated, autoinhibited ‘off’ state (K_D,actin_>100 μM; ATPase activity >1000-fold reduced) ^18,19^and an enzymatically active ‘on’ state is critical for cell function^18^. Disruption of this equilibrium causes changes in cytoskeletal dynamics and mechanical properties in fundamental processes including cell adhesion and migration with implications for disease pathogenesis^20,21^. While the binary switch model has been well established with biochemical ^18,19^ and low-resolution negative stain electron microscopy (EM) studies^22-24^, the molecular details of the autoinhibited 10S ‘off’ state of NM2 remain largely unknown, despite its potential as a target for NM2-specific therapeutics.

In this work, we investigate the structural mechanism of the 10S conformation of nonmuscle myosin-2B (NM2B). The structure adopts a highly asymmetric conformation that renders NM2B inactive through the sequestration of key elements required for actin binding and filament formation. Our results also reveal that destabilization of the coiled coil in the myosin tail confers high structural flexibility in the distal portion of the folded molecule that is expected to play a role in phosphorylation-dependent autoinhibition relief and filament formation. These principles are likely to universally apply to all myosins forming the 10S conformation.

## Results

### The overall architecture of 10S

To define the structural basis of the autoinhibition mechanism of 10S, we produced full-length human NM2B in the baculovirus/*Sf*9 insect cell system and determined its structure using single particle cryo-electron microscopy (cryo-EM) analysis (Fig. 1, Supplementary Movie 1). We adopted a focused refinement strategy and locally masked the compact portion of the molecule that includes both heads and the proximal tail (∼431 Å, Supplementary Fig. 1) as well as the distal tail portion (∼104 Å, Supplementary Fig. 1) for independent data processing. Finally, the composite maps were merged to reveal the architecture of the NM2B in the 10S state at a global resolution of ∼5.24 Å (Fig. 1, Supplementary Fig. 1 h).

**Fig. 1:**
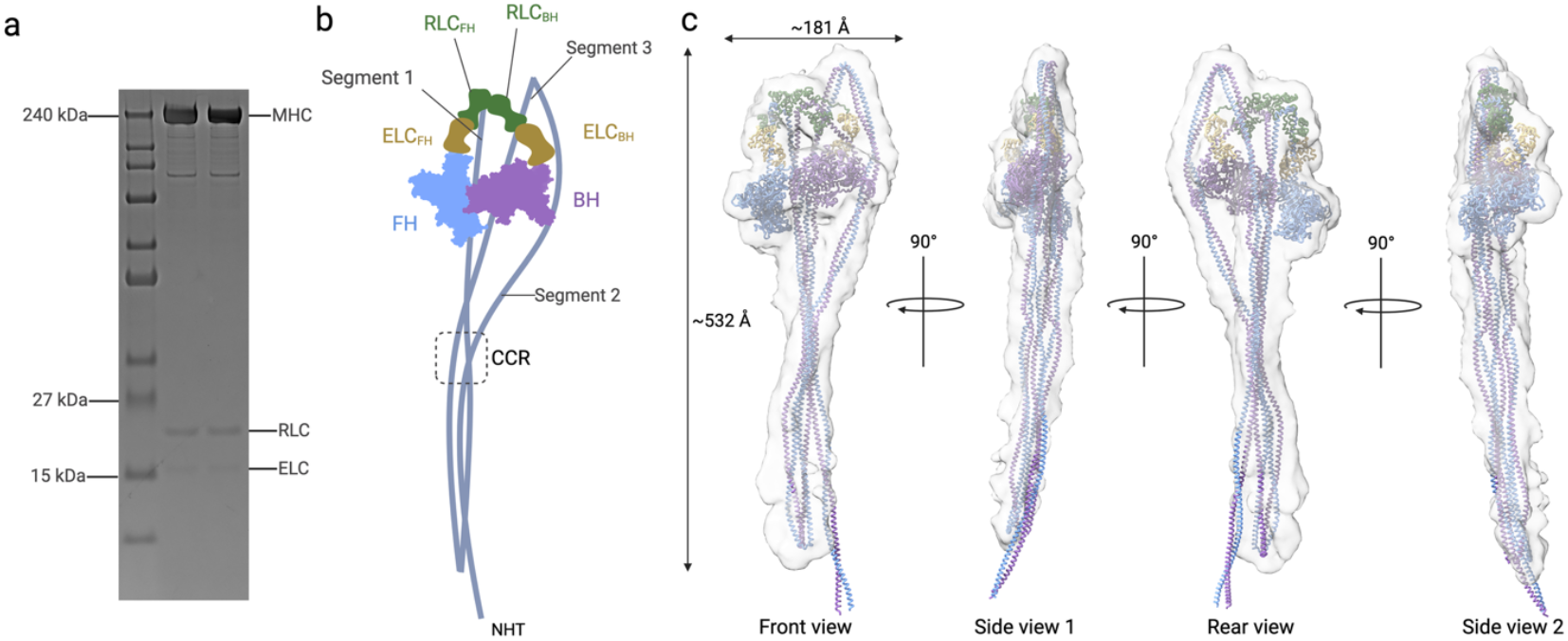
Cryo-EM structure of full-length human NM2B in the 10S conformation. **a**, SDS-PAGE gel shows full-length human NM2B heavy chain (MHC), essential light chain (ELC), and regulatory light chain (RLC). HC apparent molecular weight is ∼229 kDa, ELC is ∼16.9 kDa, and RLC is ∼19.7 kDa. **b**, Schematic representation of the autoinhibited 10S conformation of NM2B. The myosin heavy chains are designated according to whether they belong to the BH (blue) or the FH (purple). The corresponding ELCs are labeled as ELC_BH_ for the BH and ELC_FH_ for the FH. The RLCs are designated as RLC_BH_ and RLC_FH_, respectively. Tail segments, NHT, and CCR are indicated. Schematic not drawn to scale. **c**, Cryo-EM reconstruction of full-length NM2B in the 10S conformation. Various views of 10S show the overall architecture of the protein complex. The NHT is not resolved in the reconstruction.

The overall architecture of the solved 10S structure agrees with existing low-resolution negative stain EM 2D class averages of nonmuscle and smooth muscle myosin-2^22-24^. 10S is a tightly folded and elongated molecule with a length of ∼ 532 Å and a width of ∼ 181Å (Fig. 1, Supplementary Fig. 1h). The two motor domains are docked against one another and interact with portions of the folded tail in a conformation termed the interacting heads motif (IHM)^25,26^. After emanating from the structurally asymmetric motor domains, the two myosin heavy chains follow overall separate paths until they converge at the proximal end of the tail domains to form the start of the coiled coil. From this point, the two myosin heavy chains in the coiled coil tail domain follow an overall similar path that features two hairpin turns that fold the tail into a tri-segmented structure. This arrangement produces three segments, designated as segment 1, segment 2, and segment 3, that enable defined contacts with the motor domains in the IHM (Figs. 1, 2a). The two heavy chains diverge at the very distal end of the molecule and terminate in the non-helical tailpiece (NHT) (Fig. 1).

**Fig. 2:**
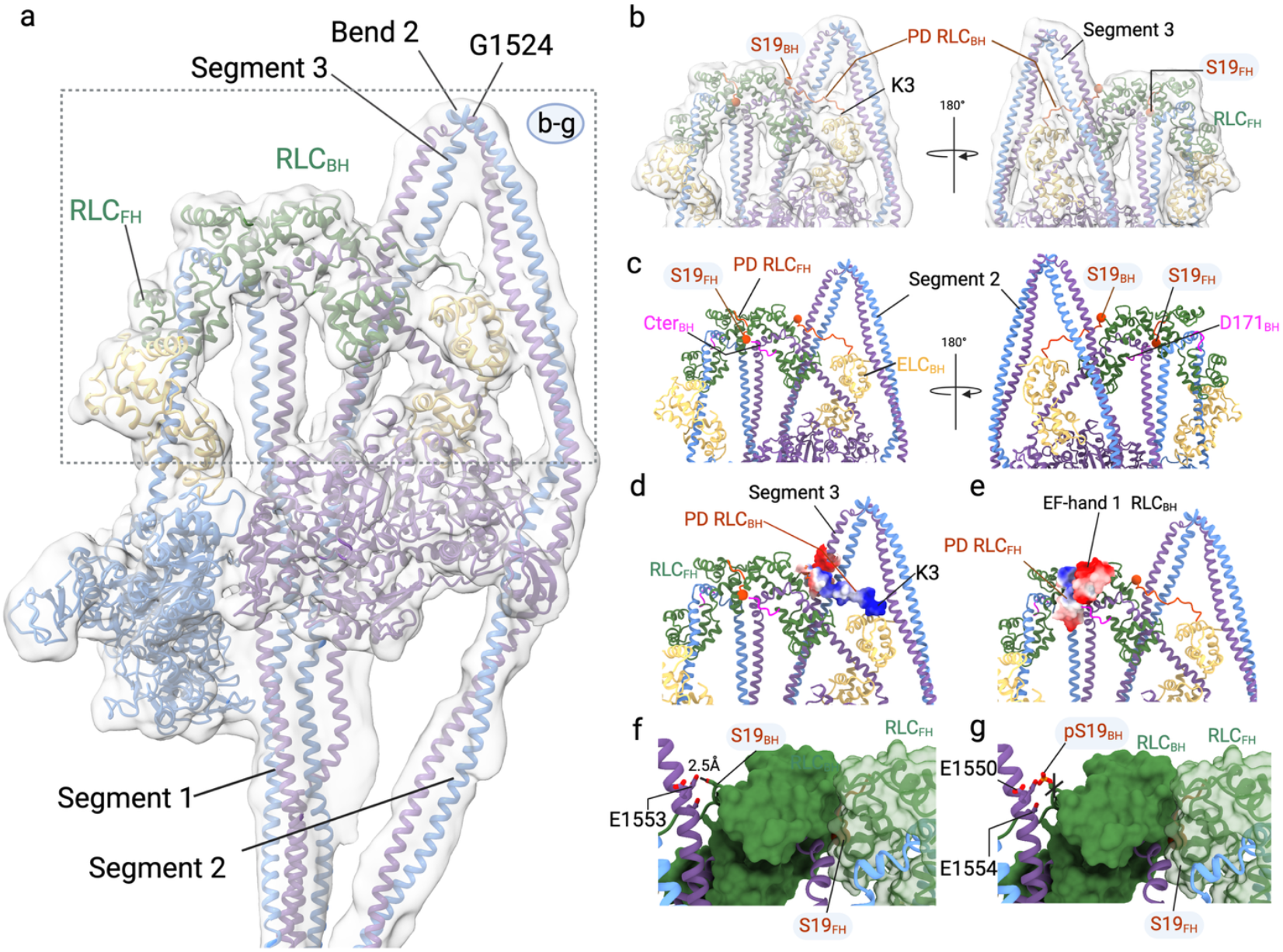
Role of RLCs in stabilizing the 10S conformation. **a**, Overview of the IHM within the 5.24 Å cryo-EM map. Key regions are indicated. **b**, The N-terminal phosphorylation domains (PD) of RLC_BH_ and RLC_FH_ are shown in orange. The PD residues K3 to T18 of the RLC_BH_ projects towards segment 3. **c**, The C-terminus of RLC_BH_ in magenta (Cter RLC_BH_) interfaces the hook region of the BH. S19 of RLC_FH_ is sandwiched between RLC_BH_ and RLC_FH_. S19 of the RLC_BH_ is in proximity to segment 3. The last residue, D171 in the Cter of the RLC_BH_ is indicated. **d**, The electropositive PD RLC_BH_ interfaces with electronegative residues in segment 3. The PD of the RLC_BH_ region is resolved until K3. That is located in close proximity to the N-terminus of ELC_BH_. **e**, The PD RLC_FH_ containing S19 residue is electropositive, and the EF-hand region in RLC_BH_ in proximity is electronegative. The orange spheres mark the position of S19 in the PD (orange) of the RLC in all panels. **f**, 180° rotated view of panel **d** showing the close-up view of S19 of the RLC_BH_ in proximity to segment 3 residues E1553, E1550, and E1554. S19_BH_ forms a hydrogen bond with E1553. S19_FH_ is sandwiched between RLC_BH_ and RLC_FH_. Select residues are shown in stick representation. **g**, 180° rotated view of panel **d** showing the close-up view of pS19 of the RLC_BH_ in proximity to segment 3 residues E1553, E1550, and E1554. pS19_BH_ clashes with E1553 (< 1Å between the residues). Select residues are shown in stick representation. The phosphoryl group is modeled onto S19 _BH_.

### Asymmetric interactions stabilize the 10S conformation and enable a sequential activation mechanism

The asymmetric arrangement of the two identical heavy chains and their associated light chains in 10S implies that individual subunits have distinct roles in stabilizing the autoinhibited state. Compared to existing crystal structures of myosin-2 motor domains in the ADP.P_i_ bound pre-powerstroke (PPS) ‘on’ state, both heads in the IHM, hereafter referred to as the blocked head (BH) and the free head (FH) ^26^have ADP and P_i_ bound in the nucleotide binding pocket but adopt distinct conformations that are incompatible with actin activation of its ATPase activity (Supplementary Fig. 2a). The motor domain cores deviate with an average root mean square deviation (r.m.s.d) of ∼1.5 Å. The lever arms are positioned ‘up’ and exhibit a pronounced kink at the pliant region between the motor domain and the ELC. The relative angles of the lever arms deviate by ∼35° between BH and FH and by ∼62° and ∼27° compared to PPS structures (Supplementary Fig. 2a-c, Supplementary Movie 2).

10S-specific intersubunit interfaces are established between the heads and their respective light chains within the IHM. The buried surface area between the BH and its associated light chains is ∼4,673 Å^2^, while the corresponding interface between the FH and its associated light chains is ∼3,617 Å^2^. Another interface is established between the ELC_BH_ and loop-1_BH_ (Supplementary Fig. 2c). Molecular interactions are also present between loop-1_FH_, ELC_FH_, and segment 1, whereas the RLC_FH_ does not interface with the motor domain (Fig. 1, Supplementary Fig. 2c,d). Direct comparison of RLC_BH_ and RLC_FH_ reveals major conformational differences (RMSD = 4.2 Å), including a rotation around the flexible linker that positions their N- and C-termini in opposite directions (Supplementary Fig. 2d, e). Interactions between the light chains and the motor domain causes pronounced differences in the pliant and hook conformations of the BH and FH myosin heavy chains (Supplementary Fig. 2). The comparison of the RLCs in the IHM to those in a published PPS structure (PDB ID: 1QVI) showed that the RLC_FH_ aligns more closely with RLC_PPS_ (RMSD = 3.5 Å) than with RLC_BH_ (RMSD = 7.4 Å) (Supplementary Fig. 2e).

A new stabilizing interface between the two heavy chains is formed between the RLC_BH_ and the RLC_FH_, which is mediated by the phosphorylation domain (PD_FH_) of the RLC_FH_ (Fig. 2) that is resolved until residue K3. The PD_BH_ runs across segment 3 and interfaces with the N-terminus of ELC_BH_ (Fig. 2c). The phosphorylatable residue S19_FH_ in the PD_FH_ is part of the interface and is buried between both RLCs rendering it inaccessible for phosphorylation by RLC kinases (Fig. 2b, c). This autoinhibitory ‘lock-and-key’ module, primarily formed by the RLCs at the apex of the 10S structure, holds both heavy chains together. The positively charged PD_BH_ which includes the phosphorylatable S19_BH_ emanates from the RLC_BH_ and contacts a negatively charged surface in segment 3 (Fig. 2d,e). Similarly, the negatively charged EF-hand 1 region of the RLC_BH_ interacts with the positively charged N-terminus of the RLC_FH_ (Fig. 2e). Together, the unresolved PD regions and electrostatically complementary interactions contribute to the stability of 10S in the ‘off’ state (Fig. 2d,e). The different positions of the RLC PDs in 10S suggest that 10S relief through S19 phosphorylation occurs first in the accessible PD of the RLC_BH_ (Fig. 2f,g). The introduction of a negative charge by phosphorylation is likely to disrupt favorable electrostatic interactions with the negatively charged region in segment 3 and to trigger structural transitions in the RLC_BH_ that lead to the destabilization of the autoinhibitory module (Fig. 2f,g). Subsequent conformational transitions in the RLC_FH_ likely expose the PD_FH_ and allow S19_FH_ phosphorylation and the transition into the 6S conformation.

### Mechanism underlying the reduced actin affinity of 10S

Actin serves as a nucleotide exchange factor for myosin and its binding to the motor domain strongly activates the ATPase activity of myosins in the ‘on’ state^7,27^. The actin binding cleft between the U50 and the L50 is wide open (> 27 Å) in both motor domains compared to known rigor or ADP bound structures (Fig. 3). Major structural elements involved in actin binding (Fig. 3) are sequestered to stabilize the IHM, thereby preventing the force-generating interaction with actin. In most crystal structures of myosins in the ‘on’ state, the major actin binding element loop-2 ^28^ is largely disordered and projects away from the head domain. In the 10S structure, loop-2_BH_ is tucked up into the actin binding cleft of the BH (Fig. 3 and Supplementary Figs. 3, 4). This sterically hinders positively charged residues in loop-2 that are critical for initiating actin binding and promoting cleft closure and prevents the transition of the motor domain from a weak into the strong actin binding state^29-31^. Loop-2_BH_ also shields the helix-loop-helix (HLH) motif from making strong hydrophobic and electrostatic interactions with actin. Loop-2_FH_ and the cardiomyopathy loop_FH_ (CM-loop) interact with segment 1_FH_ (Supplementary Fig. 3c, d). The CM-loop_BH_ and loop-2_BH_ stabilize the BH:FH interface (Supplementary Fig. 3a, b). Loop-2_FH_ and HLH_FH_ are oriented towards the surface of the FH, but segment 1_FH_ sterically hinders the formation of a strong actin binding interface (Fig. 3 and Supplementary Figs. 3,4). In addition, inter-head and head-tail interactions cause an increase in the overall negatively charged IHM surface, which further disrupts complementary charge interactions with the negatively charged actin filament (Supplementary Fig. 5, Supplementary Movie 3). Collectively, the sequestration of actin binding elements within the IHM together with the overall negative charge of 10S explains the markedly reduced actin affinity when compared to myosins in the ‘on’state^18,19^.

**Fig. 3:**
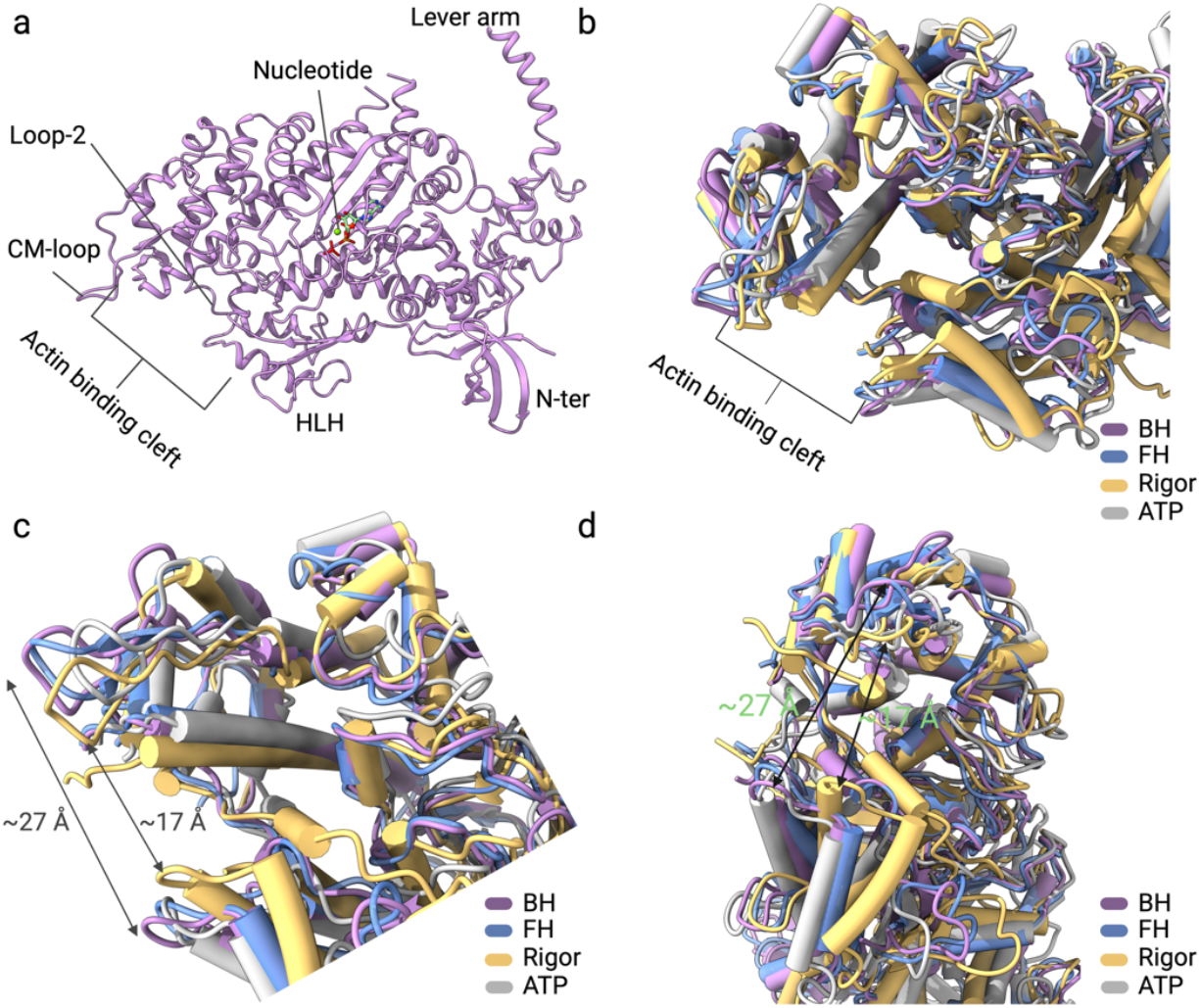
Open conformation of the actin binding cleft in NM2B 10S. **a**, Structure of the NM2B motor domain with key structural elements indicated. **b**, Superimposition of the motor domains of the BH (purple), FH (blue), rigor (PDB ID: 5JLH, gold), and post-rigor (ATP-bound, PDB ID: 1MMN, grey) structures. **c–d**, Close-up views of the superimposed actin binding regions of the myosin motor domains in the BH (purple), FH (blue), rigor (PDB ID: 5JLH, gold), and post-rigor (ATP-bound, PDB ID: 1MMN, grey) structure. Cleft widths are shown for myosin structures with high (rigor) and low (BH, FH, and ATP-bound) actin affinity. The width of the actin binding cleft, measured between residues of the CM-loop and the helix-loop-helix region (HLH), is indicated.

### Complex folding of the tail

The long coiled coil tail makes up two-thirds of the length of the extended 6S monomer (Figs. 1, 2) and is essential for switching ‘off’ NM2 activity in the autoinhibited 10 conformation through a complex folding mechanism^32^. However, the structural mechanisms by which the tail folds and interacts with itself distal to the IHM remain largely unknown. Our structure enabled us to define the molecular architecture of the tail segments that are too closely juxtaposed to be discerned by negative stain electron microscopy^22,23,32^ or cryo-EM on smooth muscle myosin-2, where >350 Å of the folded tail remained unresolved^33-35^. The NM2B 10S structure (Fig. 4, Supplementary Figs. 1b-d, 6) shows that the folding of the tail results in three segments of ∼ 460 to 530 Å length^23^. Acute folding of segment 1 at the distal end of 10S (bend 1) gives rise to segment 2 and creates an initially antiparallel arrangement of segments 1 and 2. Segment 2 diverges from this antiparallel interaction with a U-shaped bend formed by a stretch of residues D1160 to Q1166 and then runs across and beyond the BH in the IMH. It then bends between residues D1522 to N1526 in bend 2 to give rise to segment 3, which lines the BH and runs as a relatively straight segment towards the distal end of 10S, where it extends beyond bend 1 (Figs. 1, 4) and terminates in the NHT. The tail length, measured between amino acids K945 at the distal end of the IHM and A1164 at bend 1 is ∼325 Å. The tip-to-bottom length is ∼532Å and the overall contour length of 10S is ∼54 nm. This is in good agreement with previous measurements (∼50.1 nm) obtained from negative stain EM studies^22^.

**Fig. 4:**
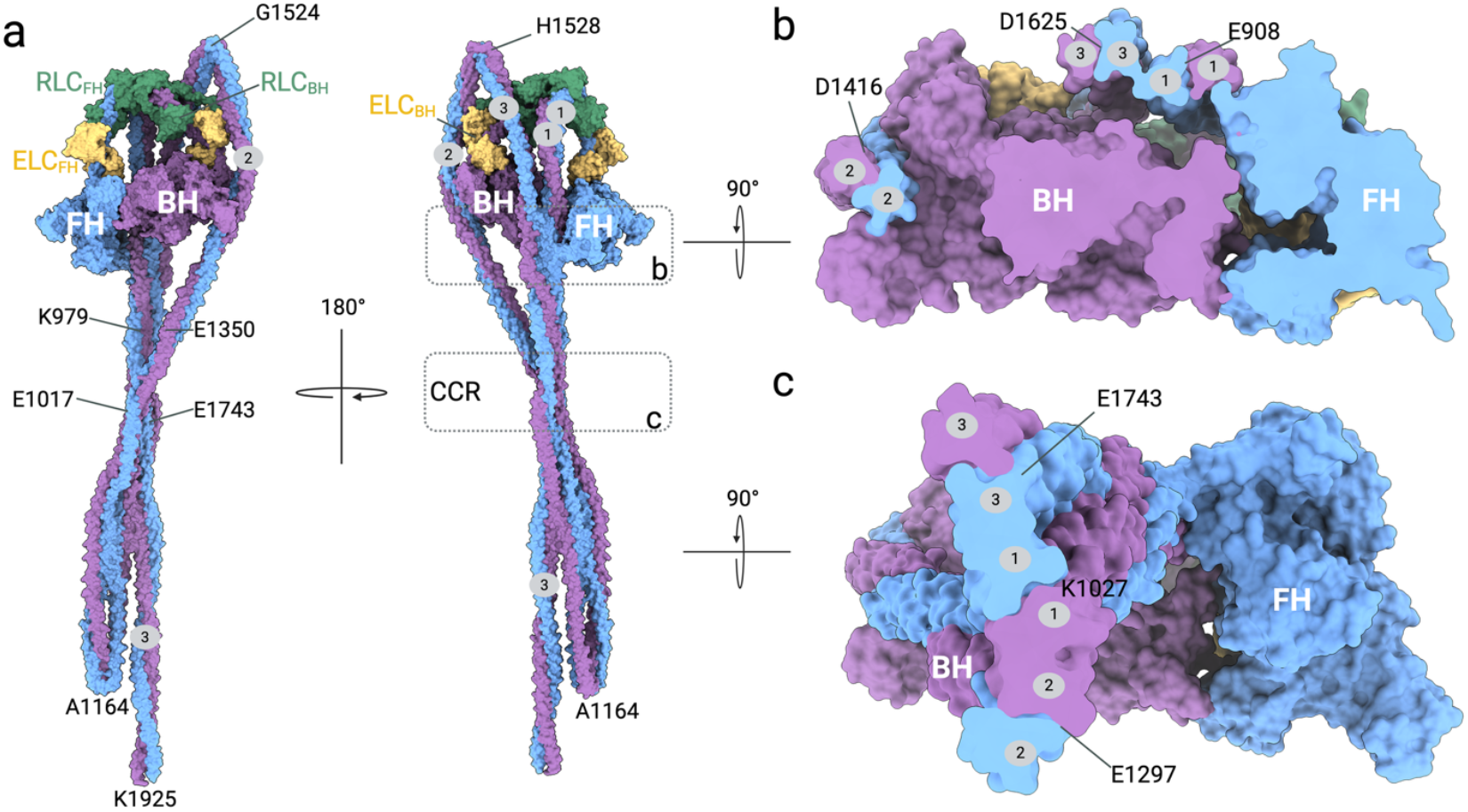
Intersubunit interfaces in the 10S conformation. **a**, Front and rear views of the NM2B 10S structure. The FH is colored blue, the BH is colored purple, and the corresponding ELC and RLC for each heavy chain are highlighted in green and yellow, respectively. Tail segments 1, 2, and 3 are numbered, and the CCR and selected key residues are annotated. **b**, Close-up cross-sectional view of the BH and FH regions, highlighting the spatial organization of the proximal tail segments. Tail segments 1, 2, and 3 are numbered, and the CCR and selected residues are annotated. **c**, Close-up view of the proximal tail region, highlighting the convergence of segments 1, 2, and 3 in the CCR.

The three tail segments converge at a central crossover region (CCR) (Fig. 4, and Supplementary Fig. 5, Supplementary Movies 3, 4) where the tail segments are arranged side-by-side, with segment 1 sandwiched between segments 2 and 3 (Figs. 1,4, and Supplementary Movies 3, 4). This ‘multi-lock’ folding at the CCR is stabilized through electrostatic interactions. The CCR overlaps with the assembly competence domain (ACD, residues 1729-1763) 2 in segment 3, while the portion of segment 3 distal to the CCR contains the ACD1 (residues 1875-1913) and the NHT that are critical for the assembly of NM2 into filaments in the ‘on’ state (Supplementary Fig. 7a, Supplementary Movie 5)^36-38^.

### Flexibility of the tail in the autoinhibited conformation

The presence of a ∼ 1,500 Å coiled coil in the extended myosin tail results in an alternating pattern of positive and negative charged zones along the length of the tail that has been implicated in the parallel, symmetric alignment of two myosin heavy chains in the 6S monomer and the antiparallel alignment of adjacent tail domains in a myosin-2 filament^39-41^. Three skip residues (Q1193, Q1586, K1811), that are not involved in the regular heptad repeat pattern characteristic of coiled coils, are believed to introduce flexibility and maintain the periodic charge distribution (Supplementary Figs. 7,8, and Supplementary Movie 5)^41,42^. The skip residues do not coincide with bend 1 or bend 2 (Fig. 4) in 10S, suggesting that structural alterations in adjacent regions of the coiled coil are necessary to accommodate the bending of the myosin heavy chains. This observation implies that the coiled coil in 10S actively accommodates and transmits large conformational rearrangements that necessitate deviations from the canonical coiled coil geometry characteristic of the extended molecule.

During 3D classification, we identified a distinct class exhibiting altered segment orientations compared to our consensus full-length cryo-EM reconstruction (Figs. 4 and 5). Analysis of this map revealed that segment 3 bends at the CCR and deviates from its typical overlap with segments 1 and 2 (Figs. 4 and 5a,b). In addition, segments 1 and 2 undergo conformational changes and can reposition relative to those observed in the full-length reconstruction (Figs. 4 and 5b,c).

**Fig. 5:**
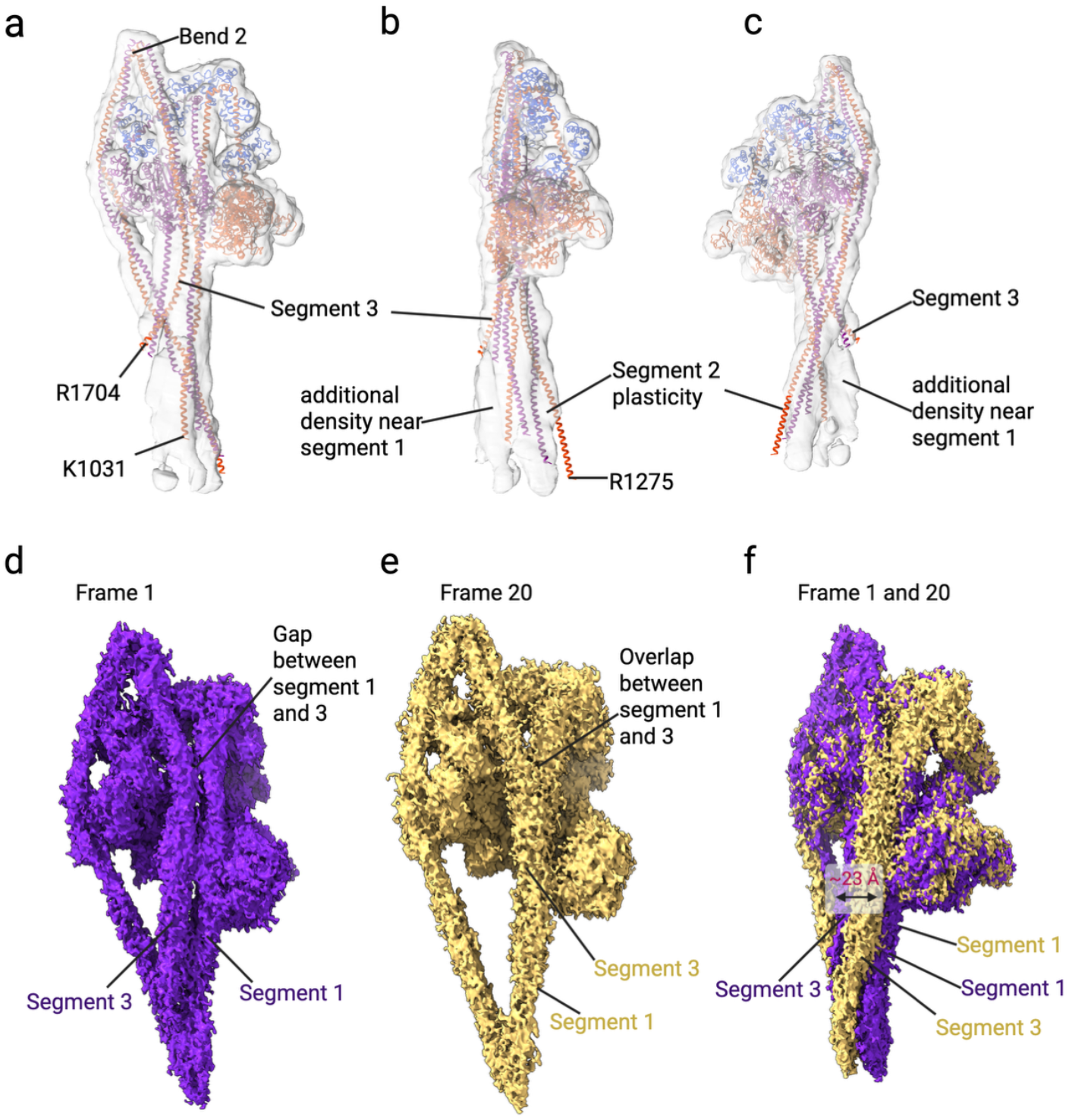
Conformational plasticity in 10S structure. **a**, Cryo-EM map showing altered densities at the CCR compared to our full-length map. Segment 3 shows an additional density ∼ 236 Å away from bend 2, and it disengages from segment 1. **b-c**, A 90° and 180° clockwise rotation of the panel **d** map shows that segments 1 and 2 adopt alternate conformations at the CCR. For clarity, only one region with segments 1 and 2 was modeled, while additional density indicates that alternative conformations are possible for these segments. **d**, Frame 1 of the principal component analysis shows that segment 3 is positioned adjacent to segment 1. **e**, Frame 20 of the principal component analysis shows that segment 3 is atop segment 1 to form the CCR. **f**, Superimposition of frames 1 and 20 of the principal component analysis shows the conformational plasticity in 10S.

To further gain mechanistic insights into the conformational plasticity of 10S, we performed 3D variance analysis (3DVA). Analysis along the leading principal component yielded 20 frames that exhibited pronounced movements of segments 1 and 3 (Fig. 5d-f). In frame 1, segment 3 is positioned adjacent to segment 1 (Fig. 5d). By frame 20, segment 3 shifts atop segment 1 to form the CCR (Fig. 5e). The 3DVA analysis further showed that the movement of segment 3 results in the formation of distinct interfaces with segment 1 (Fig. 5f, Supplementary Movie 6) in the 10S conformation. Notably, major conformational changes during the 3DVA analysis are restricted to the tail segments, whereas the heads remain relatively stable throughout the principal component trajectory. Of the two heads, the BH displays reduced motion compared to the FH, due to extensive stabilizing interactions that limits its conformational flexibility. This observation underlines that disruptions of the canonical coiled coil packing contribute to the inherent flexibility in the folded tail of 10S.

### Disease-associated mutations in 10S

NM2B is highly missense constrained^43^ and thus disease-associated mutations (Fig. 6, Supplementary Table 2) are rare but likely have pronounced effects on myosin structure and function. While some missense mutations can be mechanistically interpreted through primary sequence analysis, mutations within the myosin motor domain (W37C, D74N, Y255C, R270C, G313V, N312S, L385I, Q484E, G648D, G703D, E706K, R709Q, G743S) may alter the enzymatic properties of NM2B in its ‘on’ state. Additionally, mutations located at 10S intersubunit interfaces have the potential to disrupt the ability of NM2B to adopt the autoinhibited conformation or affect its stability. The latter ones comprise mutations R270C, N312S, G313V, L385I, and G648D in the BH-FH interface in the BH myosin heavy chain (Fig. 6). In addition, a cluster of three mutations (R1704Q, E1707K, and E1709G) is located near the CCR (Fig. 6), while mutation E1280Q is located distal to the CCR (Figs. 4,6). These mutations have the potential to influence myosin function through altered 10S stability with direct consequences for the 10S-filament equilibrium and thus cellular force generation. Mutation K1809R is located near the skip 3 residue K1811 and may impact the bending propensity of the myosin tail. Mutations within the free head motor domain near the actin-binding interface, and in segments 1 and 2 of the FH, may influence enzymatic kinetics and structural stability, but are not expected to disrupt the folding of segment 3 (Fig. 6B). Nonsense mutations R836* and E908* introduce a premature stop codon and thus result in a short myosin fragment that is incompetent in adopting the 10S conformation, irrespective of the location of the mutation in the BH or FH heavy chain.

**Fig. 6:**
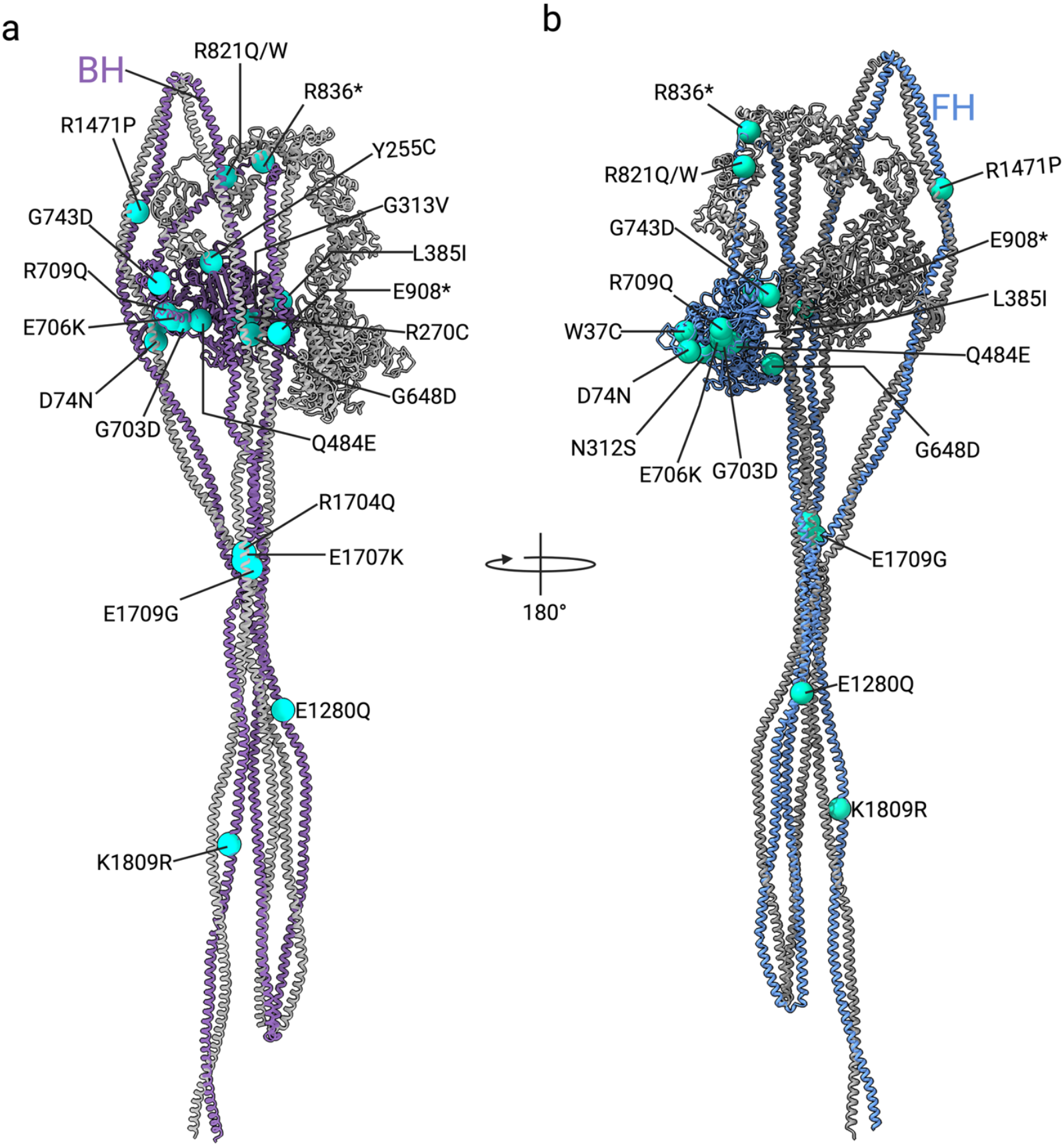
Positions of disease-associated mutations in the 10S structure. **a**, Select disease-associates mutations (cyan spheres) mapped onto the BH heavy chain colored in dark violet. A cluster of mutations (E1709G, R1704Q, E1707K) is located in the CCR. Mutation K1809R is located near skip residue K1811 at the distal end of the tail domain. * indicates termination. **b**, Select disease-associates mutations (green spheres) mapped onto the FH heavy chain colored in blue.

### Comparison between IHM in class-2 myosins

The IHM is widely conserved throughout evolution as an ancient mechanism to suppress the enzymatic activity of myosin-2s ^44^ and has been observed in isolated myosin molecules^23,33-35,45,46^, isolated muscle fibersb^47^, and living organisms^48^. Comparison of the NM2B 10S structure with that of cardiac myosin-2 in the autoinhibited state (PDB ID: 8Q6T) shows a significant deviation in the positioning of FH, the light chains, and segment 1 (Fig. 7, Supplementary Movie 7). Cardiac myosin-2 shows reduced interdomain interactions and lacks stabilizing interactions with segments 2 and 3, contributing to its reduced IHM stability^49^ compared to smooth and cardiac myosin-2 (Fig. 7a, b, Supplementary Movie 7) as its extended tail is stabilized within the thick filament by specific partner proteins in the autoinhibited conformation. Superimposition further shows a pronounced difference in axial orientation between segment 1 of the two structures as it emanates from the IHMs of cardiac myosin-2 and NM2 (Fig. 7c). Whereas the segment 1 in NM2B lies relatively flat, the corresponding segment in cardiac myosin-2 exhibits a ∼25° upward tilt. Segment 1 of cardiac myosin-2 shows a displacement of 40 Å in the IHM region, > 50 Å at the CCR, and > 121 Å at the distal region of segment 1. The BH of NM2 aligns more closely with that of cardiac myosin-2 than the FH and segment 1. shows a better overlap with cardiac myosin-2 compared to the FH and segment 1.

**Fig. 7:**
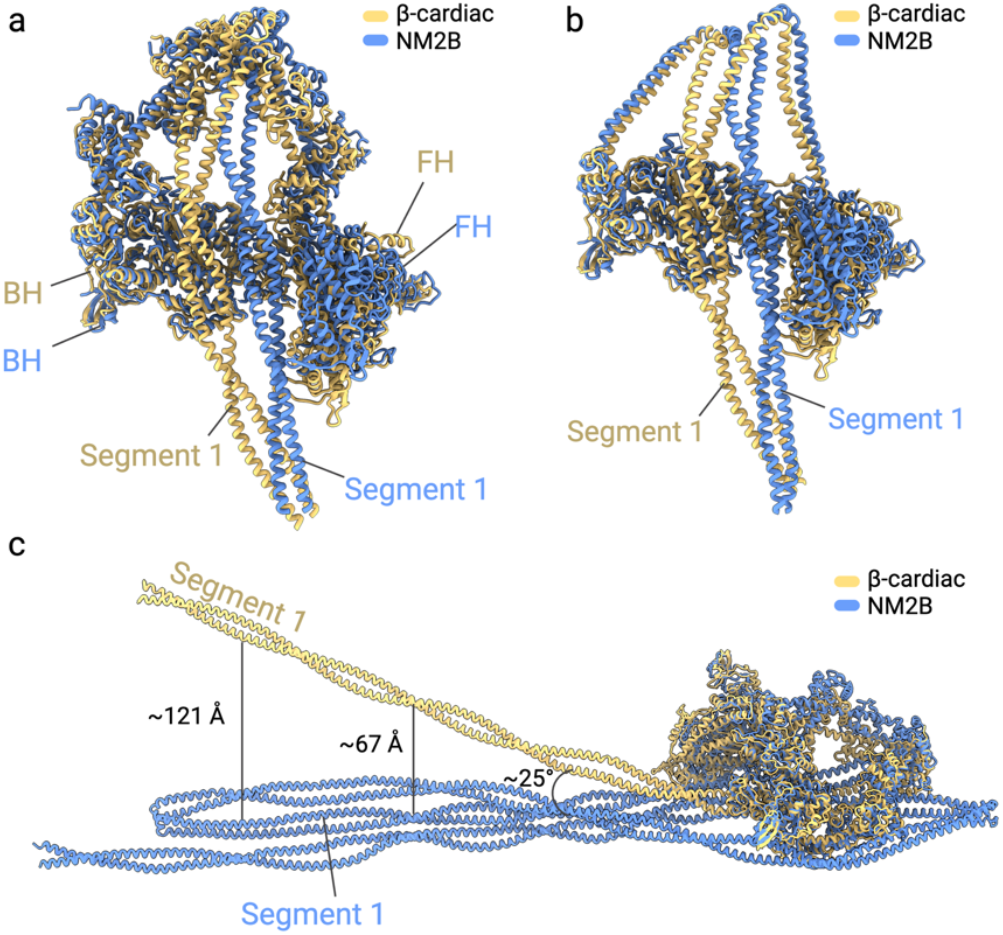
Structural differences in the autoinhibited states of class-2 myosins. **a**, Superimposition of the IHM portion of NM2B (blue) in the 10S conformation and the IHM of β-cardiac myosin-2 (PDB ID: 8Q6T, yellow). **b**, Close-up view of IHM structures of NM2B (blue) and β-cardiac myosin-2 (PDB ID: 8Q6T, yellow). The BHs of NM2B and β-cardiac myosin-2 show significant structural overlap compared to the FHs. Light chains are not shown for clarity. **c**, Segment 1 of NM2B (blue) deviates >25° compared to β-cardiac myosin-2 (yellow). When superimposed with the BH, segment 1 shows a displacement of 67 Å in the CCR, and > 120 Å at the distal region of segment 1.

## Discussion

Recent cryo-EM studies have elucidated the IHM structures of smooth and cardiac myosin-2s^33,35,46^, but the molecular architecture of the IHM of NM2 and the tail path beyond the IHM remains largely uncharacterized. Using single-particle cryo-EM, we determined the structure of NM2B in its autoinhibited 10S conformation. Complex intersubunit interactions between the BH and the FH form asymmetric interfaces that effectively turn ‘off’ myosin motor activity (Fig. 4). Our structure shows that the two identical myosin heavy chains adopt distinct conformations (Fig. 1, 4) that result in the sequestration of actin binding elements as stabilizers of inter- and intrasubunit contacts to collectively stabilize the 10S conformation and to suppress actin activation of the myosin ATPase activity (Fig. 3, Supplementary Fig. 3,4). Additional stabilizing interfaces are formed between the motor domain and the ELC, the ELC and the RLC, and both RLCs (Fig. 1, 2, 4a). Distinct stabilizing interactions between the light chains bound to the neck domain of the BH and the FH heavy chains (Fig. 4, Supplementary Fig.2d-f) not only contribute to the asymmetry within 10S but also play different roles in the ‘on’/’off’ regulation of NM2 through RLC phosphorylation on S19.

Structural analysis indicates that the phosphorylatable S19 residues on the two RLCs are in markedly different environments in the 10S state (Fig. 2). The S19_FH_ is buried between RLC_BH_ and RLC_FH_ and is apparently inaccessible to RLC kinases, whereas S19_BH_ on the RLC_BH_ is solvent-exposed and accessible ^34^. Earlier studies found that, in a high ionic strength buffer where myosin adopts the extended 6S conformation, both RLCs are phosphorylated at a single rate, implying equivalent environments for the two S19 sites ^50,51^. However, under 10S conditions, phosphorylation proceeds with biexponential kinetics with equal amplitudes, indicating two kinetically distinct environments.

Interestingly, when the actin-activated ATPase was measured as a function of the percent of RLC phosphorylation, it was found that at 50% phosphorylation, there was little activation of the ATPase activity by actin. As the phosphorylation level increased beyond this, there was a linear increase in ATPase activity with increasing phosphorylation ^50^. However, when the myosin was phosphorylated in high-salt conditions, there was a nonlinear relationship between the extent of phosphorylation and the activation of the ATPase activity. At 50% phosphorylation, the ATPase activity was 25% of the maximum obtained at 100% phosphorylation. The shape of the curve was consistent with a model where the phosphorylation of the two heads was random and where both heads had to be phosphorylated before the ATPase of either could be activated. The data obtained when phosphorylating 10S would also be consistent with the concept that both heads need to be phosphorylated before either could be activated, but in this case, presumably, the BH was preferentially phosphorylated and was not associated with activation. Notably, smooth muscle fibers demonstrated a high level of resting phosphorylation that was not associated with force or stiffness development ^52,53^.

These findings, coupled with the structural data presented here suggest that S19 of the BH is first phosphorylated, but this does not lead to activation of the ATPase activity of either of the two heads. Instead, the proposed conformational changes in the PDs of the two heads in response to the introduction of the negatively charged group in the BH make S19 of the FH accessible to the kinase (Fig. 2d), and its phosphorylation leads to a deconstruction of the IHM allowing for productive interactions with actin. This type of mechanism would allow a myosin in nonmuscle cells or in smooth muscle to be primed by the unproductive phosphorylation of S19_BH_ such that subsequent phosphorylation of the S19_FH_ would then activate the molecule.

Contrary to the classical view of coiled coils as two intertwined α-helices, our work demonstrates significant alterations in the coiled coil geometry of NM2B in the 10S conformation compared to the canonical extended conformation. Although class-2 myosin tails are defined by nearly perfect heptad repeats ^39,54^ in the extended conformation, the presence of these motifs in the primary sequence does not fully account for the structural complexities of the asymmetric 10S conformation. Based on our 3D variability analyses and flexible refinements, we propose that deviations from the canonical coiled coil geometry are essential for NM2 regulation by both stabilizing the 10S conformation and by enabling the transitions between autoinhibited and active filamentous conformations (Fig. 5). Skip residues and adjacent ‘untwisting’ regions in the 10S conformation function as molecular hinges that allow for the pronounced tail flexibility of 10S, in line with results from previous negative stain EM studies^23^. These deviations in the coiled coil geometry highlight the limitations of strictly linear coiled coil prediction methods for the analysis of asymmetric protein assemblies including the IHM of smooth muscle myosin-2 and the scallop myosin-2 rod^34,55,56^, proteasomal ATPases^57^, dynein^58^, and bicaudal-D^59^ where noncanonical coiled coils have been experimentally shown.

The transition of NM2 from the autoinhibited 10S monomer to the active filamentous state is a highly regulated process essential for its cellular functions^1-3^. Our structure reveals that the folded 10S conformation sequesters key assembly domains, including ACD2 to prevent filament formation in the ‘off’ state, while it does not prevent the formation of antiparallel 10S dimers^22,60^. Upon relief of autoinhibition, conformational changes expose the ACD2 and allows NM2 monomers to self-associate via their coiled coil tails into bipolar filaments^22,60^.

Our findings also highlight the significance of the structural asymmetry of NM2B, particularly with respect to how each heavy chain adopts a distinct functional role within the 10S confirmation. The inherent asymmetry suggests that heterozygous disease-associated or structure-inspired mutations may have different functional impacts depending on whether the mutant heavy chain occupies the BH or FH position (Fig. 6). In heterozygous individuals, random dimerization of wild-type and mutant heavy chains is expected to yield a population of 25% wild-type homodimers, 25% mutant homodimers, and 50% heterodimers. The physiological consequences of pathogenic variants will therefore depend not only on the inherent properties of the mutant homodimers but also on the effect of the mutation within heterodimeric complexes, particularly as the spatial context of the mutation may alter or preserve critical intersubunit interactions within the 10S conformation. Notably, our structural work predicts that disease-associated mutations in the BH heavy chain may have more pronounced effects on 10S formation and stability and therefore, cellular function than those in the free head heavy chain, highlighting the importance of positional context for interpreting the impact of specific NM2B mutations.

In summary, our findings offer a structural framework for understanding NM2 autoinhibition mechanisms and will inform new avenues for modulating cytoskeletal architecture in various contexts.

## Methods

No statistical methods were used to predetermine the sample size. The experiments were not randomized. The investigators were not blinded to allocation during experiments and outcome assessment.

### Recombinant protein production, purification, and quality control

C-terminally FLAG-tagged human NM2B (UniProt ID: P35580) along with the human essential (ELC, UniProt ID: P60660 (MYL6)) and regulatory (RLC, UniProt ID: O14950 (MYL12B)) light chain was produced in *Sf*9 insect cells using the baculovirus expression system. Cells were maintained and infected with recombinant viruses according to the manufacturer (Thermo Fisher Scientific). The cells were harvested by centrifugation (5,000 rpm, 4°C, 10 min) 72 hours post-infection, flash-frozen in liquid nitrogen, and stored at -80°C. For purification, the frozen cell pellet was thawed on ice and lysed in a Dounce homogenizer using a buffer containing 10 mM MOPS pH 7.0, 500 mM NaCl, 10 mM MgCl_2_, 1 mM EGTA, 2 mM ATP, 1 mM PMSF, and protease inhibitor tablets (Sigma). After centrifugation (15,000 rpm, 4°C, 35 min), the clarified supernatant was applied to a FLAG affinity column (Sigma), extensively washed, and finally eluted with the 100 μg/ml FLAG peptide (Biomatik) in buffer containing 10 mM MOPS pH 7.0, 500 mM NaCl, and 0.1 mM EGTA. The eluted fractions were pooled, dialyzed against a buffer containing 50 mM NaCl, and then centrifuged at 2,000 × g for 10 minutes to concentrate the protein. The pellet was dissolved in a storage buffer containing 500 mM NaCl and frozen at -80°C for long-term storage.

### Sample preparation and Cryo-EM data acquisition

The purified NM2B sample was diluted to 1 mg/ml in low-salt buffer containing 10 mM MOPS pH 7.0, 150 mM NaCl, 1 mM EGTA, and 2 mM MgCl_2_ and incubated with 1 mM ATP for 30 minutes, and then cross-linked with 0.1% glutaraldehyde for 1 minute. The reaction was immediately quenched with 0.1 M Tris pH 8.0 and sample homogeneity was verified via negative stain EM. For cryo-EM grid preparation, 4 μl of the cross-linked protein was applied to glow-discharged C-flat 1.2/1.3 300-mesh Au grids, incubated for 1 minute, and blotted for 4 seconds at 95% humidity before plunge-freezing in liquid ethane using a Leica EM Grid Plunger 2. The grids were stored in liquid nitrogen and initially screened on a 200 kV Tecnai TF20 equipped with a K2 camera (FEI) to assess sample stability and particle distribution. Optimal grids were then transferred to a Titan Krios (Thermo Fisher Scientific) operating at 300 kV and fitted with a K2 direct electron detector. A total of 1412 movies were collected with a pixel size of 0.53 Å/pixel and a defocus range of −1.5 to −3.5 μm, yielding a total electron dose of 73 e^−^/Å^2^ per movie. Data collection was performed using SerialEM^61^, and the associated parameters are summarized in Table S1.

### Cryo-EM image processing and 3D reconstruction

All raw movies were aligned, drift-corrected, and dose-weighted using MotionCor2^62^. Defocus and contrast transfer function (CTF) parameters were then estimated using the Gctf wrapper in cryoSPARC^63^. Particle-picking routines in cryoSPARC were used to generate templates, and a small subset of reference-free two-dimensional (2D) particle classes was selected and subjected to several rounds of iterative 2D classification to remove poor-quality particles. Ten well-defined 2D classes of the IHM and distal tail were subsequently used as templates for *ab initio* 3D reconstruction in cryoSPARC. Among the resulting reconstructions, a major class displayed clear IHM and distal tail region features. To improve the local resolution of this class, a soft mask was applied to exclude flexible segments distal to the IHM in Relion^64^. Nonuniform refinements were performed in cryoSPARC after optimizing per-particle defocus parameters, resulting in a global resolution of 5.24Å at a Fourier shell correlation (FSC) = 0.143. The 3D map was then postprocessed with deepEMhancer^65^. All resolutions reported here are based on the gold-standard FSC = 0.143 criterion. Detailed data collection and image processing parameters are provided in Supplementary Table 1.

### Cryo-EM model building, refinement, and validation

We combined the cryo-EM density maps of IHM and the distal tail by taking the maximum density value at each voxel within UCSF ChimeraX^66^, thereby creating a composite map suitable for model building. We used the V-shape of the IHM region and tail region as the common region for merging the maps. To model the IHM region, we used the high-resolution smooth muscle myosin-2 structure^34^ (PDB ID: 7MF3) as a template. Subsequent segments in the myosin tail were built by integrating structural predictions from AlphaFold^67^ with coiled coil prediction software such as Marcoil^68^ and Waggawagga^69^. While these programs accurately identify linear heptad repeats within a polypeptide chain, they do not fully account for the asymmetry, bends, and skip regions characteristic of the 10S conformation of NM2B. Consequently, we relied on the high-resolution structure of smooth muscle myosin-2 (PDB ID: 7MF3) as a template to guide model building at the global map resolution of ∼5.24 Å, where side chain positioning is only approximate. It is important to note that the distal tail region is flexible, thus limiting the overall resolution of the reconstructions.

After establishing the initial model, manual adjustments to the backbone and side chains were performed in Coot^70^, followed by real-space refinement in Phenix^71^. Further minimization and refinement were conducted with Namdinator^72^, which facilitates automated molecular dynamics flexible fitting into cryo-EM maps using group ADP, real-space refinements, and validation tools in Phenix in combination with the RosettaCommons software package^73^. These iterative refinements improved the overall architecture of the modeled NM2B.

## Acknowledgments

We thank the NIH MICEF center and the Center for Electron Microscopy and Analysis (CEMAS) shared resource at the Ohio State University for support and use of the facility.

## Author contributions

Conceptualization: K.C. Investigation: S.M.H., G.G., J.R.S., and K.C. Formal analysis: S.M.H. and K.C. Supervision: K.C. Validation: K.C. Visualization: S.M.H. and K.C. Writing - original draft: S.M.H. and K.C. Writing - review and editing: S.M.H., G.G., J.R.S., and K.C.

## Data and materials availability

All data needed to evaluate the conclusions in the paper are present in the paper and/or the Supplementary Materials. Atomic coordinates generated in this study have been deposited in the PDB under accession numbers XXX. The cryo-EM maps generated in this study have been deposited in the Electron Microscopy Data Bank under accession numbers XXX.

